# Identifying Antimicrobial Peptides using Word Embedding with Deep Recurrent Neural Networks

**DOI:** 10.1101/255505

**Authors:** Md-Nafiz Hamid, Iddo Friedberg

## Abstract

Antibiotic resistance constitutes a major public health crisis, and finding new sources of antimicrobial drugs is crucial to solving it. Bacteriocins, which are bacterially-produced antimicrobial peptide products, are candidates for broadening the available choices of antimicrobials. However, the discovery of new bacteriocins by genomic mining is hampered by their sequences’ low complexity and high variance, which frustrates sequence similarity-based searches. Here we use word embeddings of protein sequences to represent bacteriocins, and apply a word embedding method that accounts for amino acid order in protein sequences,to predict novel bacteriocins from protein sequences without using sequence similarity. Our method predicts, with a high probability, six yet unknown putative bacteriocins in *Lactobacillus*. Generalized, the representation of sequences with word embeddings preserving sequence order information can be applied to protein classification problems for which sequence similarity cannot be used.

## Introduction

The discovery of antibiotics ranks among the greatest achievements of modern medicine. Antibiotics have eradicated many infectious diseases and enabled many medical procedures that would have otherwise been fatal, including modern surgery, organ transplants, and immunosupressive treatments. However, due to the prevalent use of antibiotics in healthcare and agriculture, antibiotic resistant bacteria have been emerging in unprecedented scales. Each year, 23,000 people in the US alone die from infections caused by antibiotic resistant bacteria [1]. One strategy to combat antibiotic resistance is to search for antimicrobial compounds other than antibiotics, and which may not be as prone to resistance. A promising class of such compounds are the peptide-based antimicrobials known as bacteriocins [2, 3]. Bacteriocins comprise a broad spectrum of bacterial ribosomal products, and with the increased sequencing of genomes and metagenomes, we are presented with a wealth of data that also include genes encoding bacteriocins. Bacteriocins generally have a narrow killing spectrum making them attractive antimicrobials that would generate less resistance [4].

Several computational tools and databases have been developed to aid discovery and identification of bacteriocins. BAGEL [5] is a database and a homology-based search tool that includes a large number of annotated bacteriocin sequences. BACTIBASE [6] is a similar tool, which also contains predicted sequences. AntiSMASH [7] is a platform for genome mining for secondary metabolite producers, which also includes bacteriocin discovery. BOA (Bacteriocin Operon Associator) [8] identifies possible bacteriocins by searching for homologs of *context genes*: genes that are associated with the transport, immunity, regulation, and post-translational modification of bacteriocins. RiPPquest [9] is an automated mass spectrometry based method towards finding Ribosomally synthesized and posttransationally modified peptides (RiPPs) which may include bacteriocins. Recently, MetaRiPPquest [10] improved upon RiPPquest by using high-resolution mass spectrometry, and increasing the search space for RiPPs. However, bacteriocins are hard to find using standard bioinformatics methods. The challenge in detecting bacteriocins is twofold: first, a small number of positive examples of known bacteriocin sequences, and second, bacteriocins are highly diverse in sequence, and therefore challenging to discover using standard sequence-similarity based methods, (Figure 1).

**Figure 1:**
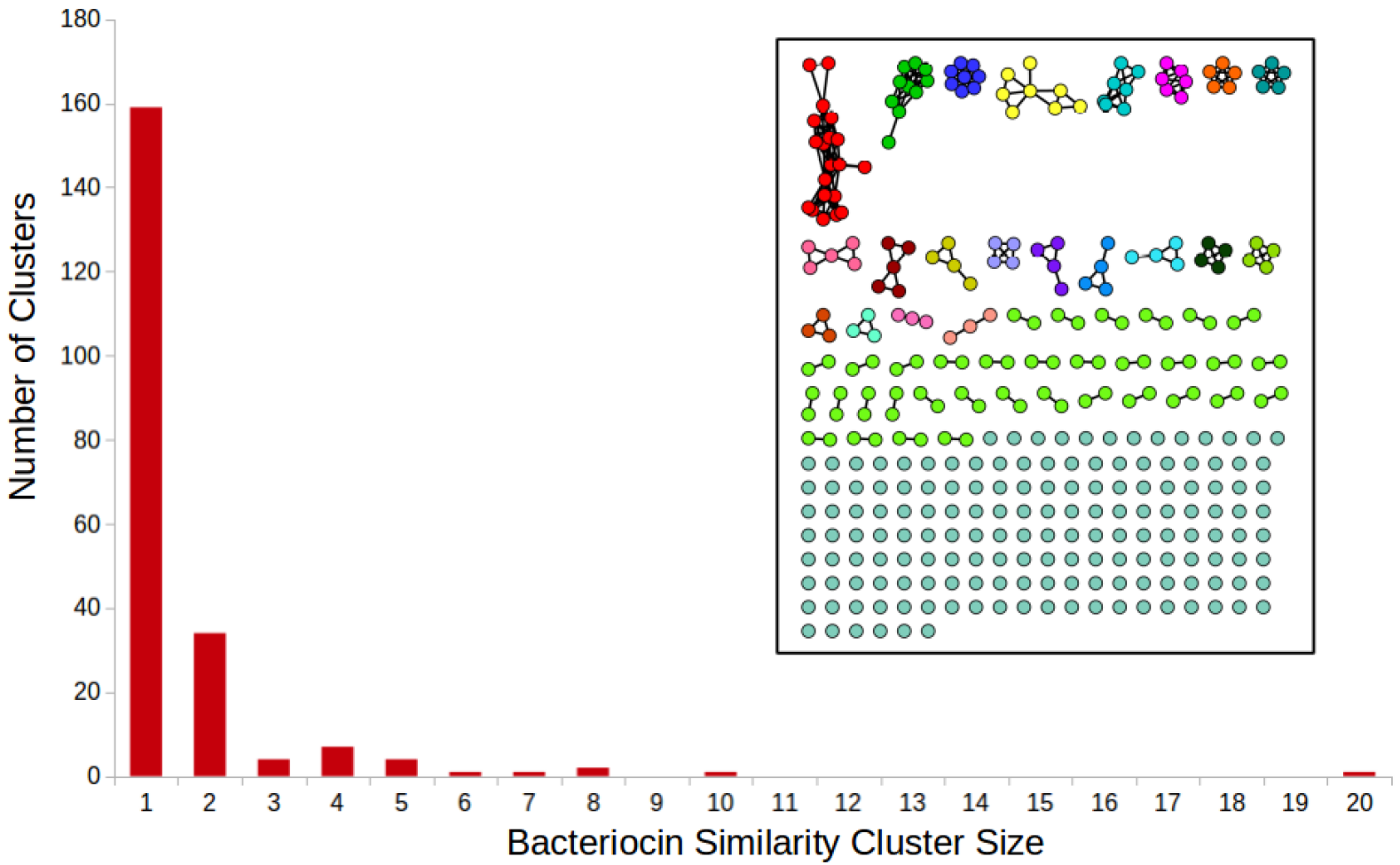
Inset: sequence similarity network for all of the bacteriocins present in the BAGEL dataset. Each node is a bacteriocin. There exists an edge between two nodes if the sequence identity between them is ≥ 35% using pairwise all-*υs*-all BLAST. The barchart shows cluster sizes.

To address these challenges we present a novel method to identify bacteriocins using word embedding. We represent protein sequences using Word2vec [11]. Using this representation, we use a deep Recurrent Neural Network (RNN) to distinguish between bacteriocin and non-bacteriocin sequences. Our results show that a word embedding representation with RNNs can classify bacteriocins better than current tools and algorithms for biological sequence classification.

## Methods

### The Representation of Proteins with Word Embedding Vectors

Word embedding is a set of techniques in natural language processing in which words from a vocabulary are represented as vectors using a large corpus of text as the input. One word embedding technique is Word2vec, where similar vector representations are assigned to words that appear in similar contexts based on word proximity as gathered from a large corpus of documents. After training on a large corpus of text, the vectors representing many words show interesting and useful contextual properties. For example, after training on a large corpus of English language documents, given vectors representing words that are countries and capitals, 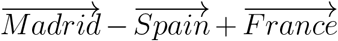 will result in a vector that is similar to 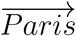, more than other vectors in the corpus [11]. This type of representation have led to better performance in downstream classification problems, including in biomedical literature classification [12], annotations [13, 14], and genomic sequence classifications [15, 16].

The training for generating the vectors can be done in two ways: the continuous bag of words (CboW) model, or the skip-gram model [11]. We adapted Word2vec for protein representation as in [17], using the skip-gram model. Instead of the common representation of protein sequences as a collection of counts of *n*-grams (also known as *k*-mers) using a 20 letter alphabet, we represent protein sequences using embeddings for each *n*-gram, covering all possible amino-acid *n*-grams (we used *n* = 3, leading to 20^3^ = 8,000 trigrams). Each trigram is a “word”, and the 8,000 words constitute the vocabulary. The Uniprot/TrEMBL database [18] constitutes the “document corpus”.

The skip-gram model is a neural network where the inputs and outputs of the network are one-hot vectors with our training instance input word and output word. A one-hot vector is a boolean vector of the size of the vocabulary (8,000 in our case, six in Figure 2b), in which only the entry corresponding to the word of choice has a value of True. We generated the training instances using a context window of size ±5, where we took a word as input and used all of its surrounding words within the context window as outputs. The process is explained in Figure 2. At the end of the training, a 200 dimensional vector for each trigram was generated by the neural network. The goal of this training was to have the 200 dimensional vectors capture information about the surroundings of each trigram that they are representing. In this fashion, we capture the contextual information for each trigram in our corpus of protein sequences. The size of the vector is a hyper-parameter which we decided upon based on the final supervised classification performance. Vectors of sizes 100, 200, and 300 were generated, and size 200 was chosen. Similarly, context window sizes of 3, 5 and 7 were tested, and size 5 was chosen.

**Figure 2:**
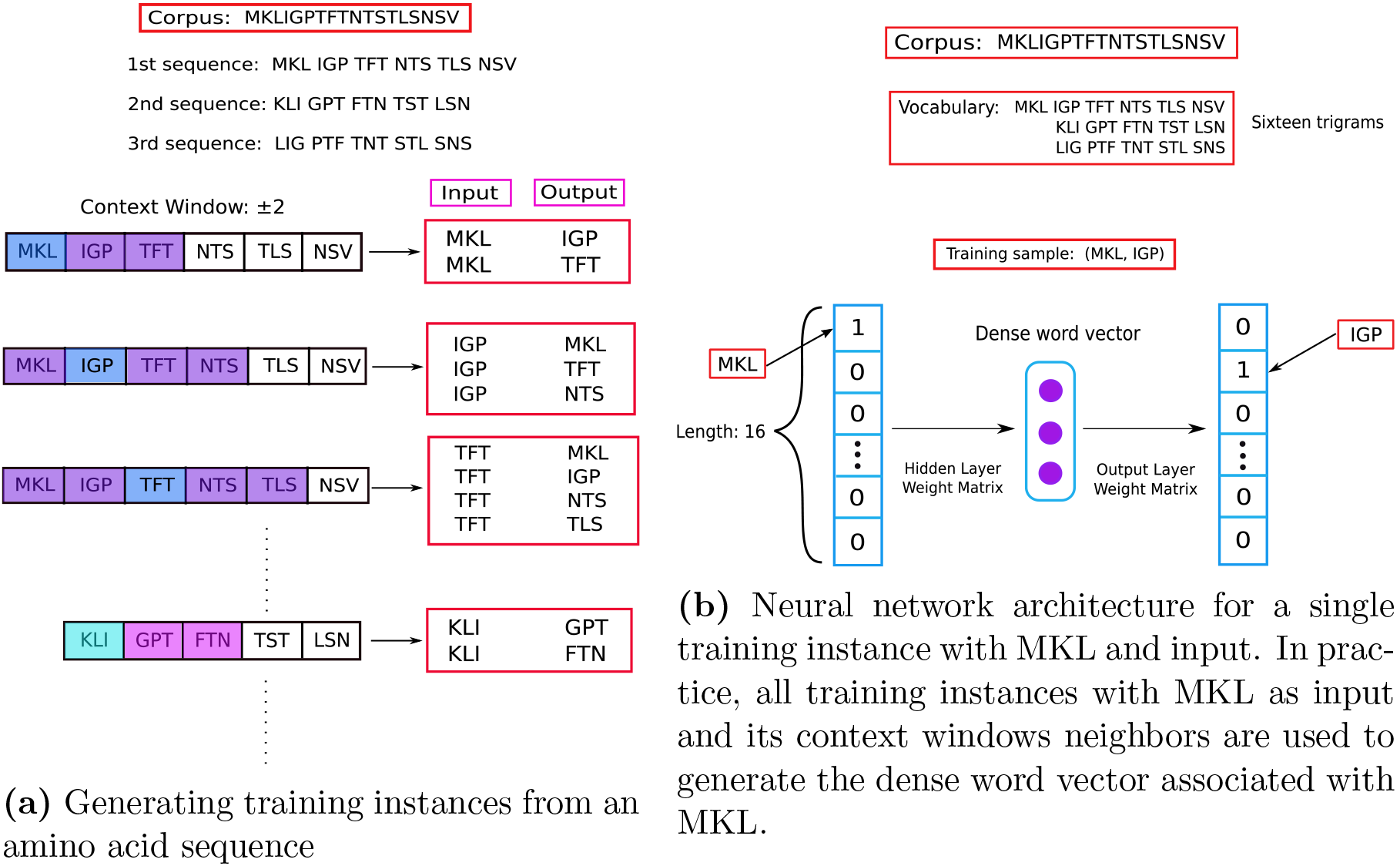
A simplified example showing representation learning for trigrams with skip-gram training. For simplicity, in the example, the vocabulary comprises of 16 words, the context window is ±2 (in our study, the vocabulary size was 8,000, and the context window ±5). **(a)** For each sequence in the TrEMBL database we created 3 sequences by starting the sequence from the first, second, and third amino acid as in [17]. This makes sure that we consider all of the overlapping trigrams for a protein sequence. A protein sequence, is then broken into trigrams, and training instances (input, output) are generated according to the size of the context window for the subsequent step of training a neural network. **(b)** The neural network architecture for training on all of the instances generated at (a). The diagram shows training on the instance where MKL is input, and IGP is output which is the first instance generated at (a). At the end of the training, for each trigram a dense word vector of size 200 is produced.

### Word2vec with a Recurrent Neural Network

We use the word-embedding representation for each trigram present in a protein sequence. We use a Recurrent neural network (RNN) to take all trigram embedding vectors as its input to represent a certain protein sequence. RNNs share the same weights for all inputs in a temporal sequence. We take advantage of this architecture by using an embedding vector of size 200 for each overlapping trigram in a protein sequence. By using the embedding vectors of overlapping trigrams as temporal inputs to an RNN, we are preserving the order of the trigrams in the protein sequence. Regarding the architecture of the RNN, we used a two-layer Bidirectional RNN with Gated Recurrent Units (GRU) to train on our data. Our hyper-parameters of number of neurons, network depth, and dropout [19] rate were determined with nested cross-validation. Since we had a small dataset, we used a dropout rate of 0.5 for the first layer, and 0.7 for the second layer. Both layers had 32 GRU units. We used a fixed number of 100 epochs for training which was also decided by nested cross-validation. For optimization, the Adam [20] method was used.

**Figure 3.**
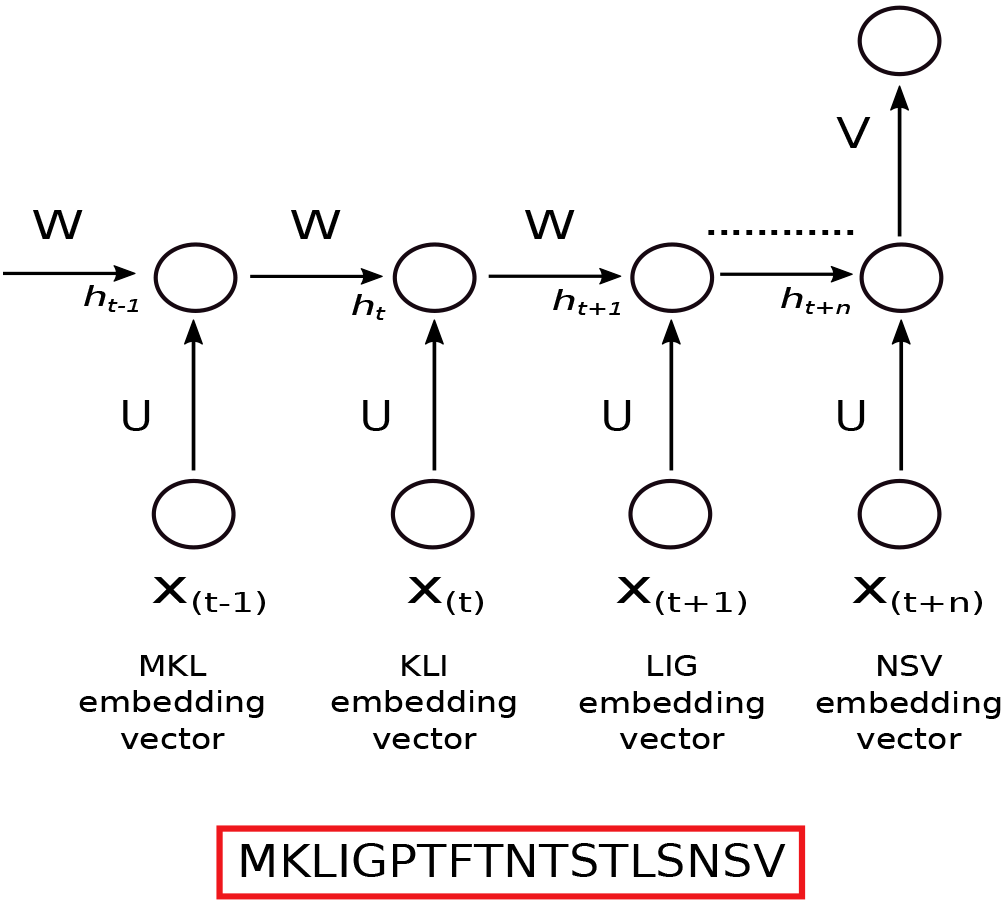
We used embedding vectors of each individual overlapping trigram present in a protein sequence as input into a Recurrent Neural Network. *X*_(*t*)_ is the input at time step *t*. In our case, at each time step *t*, input is the embedding vector of the trigram at that time step. *h*_(*t*)_ is the hidden state at time step *t*. It contains information from the previous inputs as well as the current input. This works like the memory of the network, and because of its mechanism, modern RNNs can preserve information over long ranges unlike traditional models like hidden Markov models. *U*, *V*, *W* are weights of the network. As they are being shared over all the inputs, this greatly reduces the number of parameters of the network helping towards generalization. At the end of the sequence, the network produces a prediction *y* of whether the sequence is a bacteriocin or not. In practice, we used a bidirectional RNN (not shown in figure).

### Comparing with baseline methods

We compared the performance of our method with three baseline methods: a trigram representation, BLAST, and HMMER.

We used the popular trigram representation of sequences in bioinformatics to understand the gain of accuracy, if any, using word embedding over simple trigram based representation. In this case, we created an 8,000 size vector for each sequence where the indices had counts for each occurrence of a trigram in that sequence. In this representation, the order of the trigrams are not preserved. Since the vector is sparse, we used truncated Singular Value Decomposition (SVD) to acquire the most importance features, and reduce the size of the vector. We tried with sizes of 100 and 200, and used the one that led to better classification performance. We then used these vectors with a support vector machine, logistic regression, decision Tree, and random forest, to classify between bacteriocins and non-bacteriocins.

**Figure 4:**
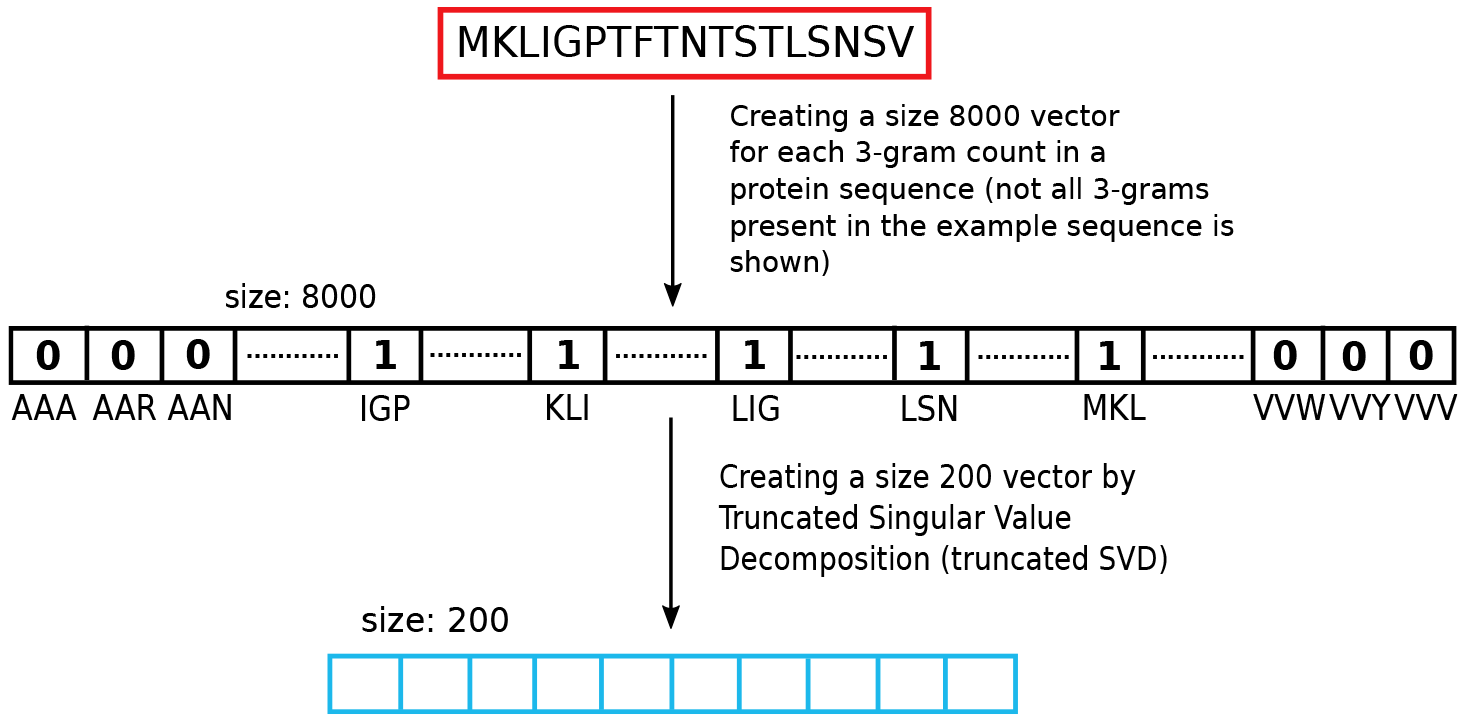
We represented each protein sequence with the overlapping trigram counts present in that sequence. This leads to a size 8,000 sparse vector. The vector was reduced to a vector of size 200 using Singular Value Decomposition. We used the size 200 vector as the baseline representation.

We also compared the performance of our method with BLAST, the method of choice for sequence similarity search. We use BLAST to see if machine learning, alignment free methods, do indeed improve performance over alignment based methods to identify potential bacteriocins. We used a 35% sequence identity score as a threshold to assign a bacteriocin label to a protein sequence.

We also compared our performance with another popular alignment based method, HMMER [21], which constructs profile hidden Markov models or pHMMs from multiple sequence alignments. In turn, the pHMMs serve as an acurate tool for sequence searching. Here we used bacteriocin pHMMs which we constructed using BOA [8]. BOA uses the BAGEL [5] dataset, and its homologs (BLAST e-value < 10^−5^) against the GenBank [22] bacterial database to build bacteriocin-specific pHMMs. We used the hmmsearch functionality provided by HMMER, and use the pHMMs from BOA to measure performance against our test set in terms of precision, recall, and *F*_1_ score.

### Building the training dataset

We used 346 sequences of length ≥ 30aa from the BAGEL database as our positive bacteriocin training samples. For the negative training set, we used sequences from the Uniprot-Swissprot [23] database. We took all the bacterial protein sequences from this database and used CD-HIT [24] with a 50% identity threshold to reduce redundancy. Then, for the primary negative training set, we took 346 sequences that had the keywords ‘not anti-microbial’, ‘not antibiotic’, ‘not in plasmid’, and that had the same length distribution as our positive bacteriocin sequences. We also generated two additional negative datasets following the same steps as above, with no overlap in the sequences between the three sets. Because identical length sequences were already exhausted by the first negative set, the length distribution of the second and third negative sets are somewhat different than the positive bacteriocin set. Figure 5 shows the length distribution of the positive, and all three negative datasets.

**Figure 5:**
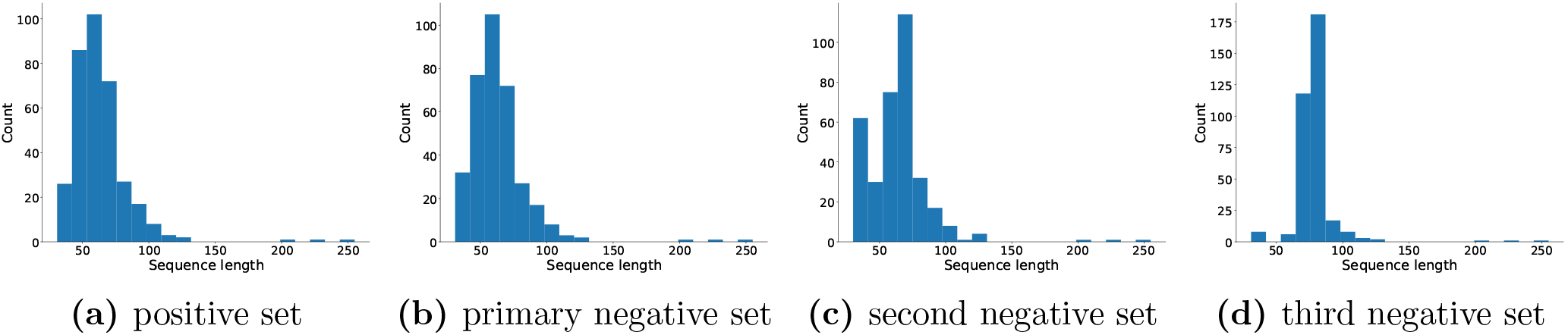
Sequence length distributions for the positive bacteriocin set, primary negative set, 2nd, and 3rd negative sets respectively. See text for details.

### Identifying genomic regions for novel putative bacteriocins

To search for genomic regions with a higher probability of containing novel bacteriocins, we took advantage of the biological knowledge of context genes which assist in the transport, modification, and regulation of bacteriocins. Many bacteriocins have some or all of four types of context genes in proximity [25, 26], (Figure 6). Having an experimentally verified set of fifty-four context genes from [8], we collected the annotation keywords for these context genes from the Refseq database, and BLASTed the BAGEL bacteriocins against the non-redundant protein database. We removed the top hits from the result which are essentially the bacteriocins themselves. We then took all the genes with similar keywords to our experimentally verified context gene set surrounding these bacteriocins within a region of ±25kb. After running CD-HIT [27] to remove redundancy, we had 1,240 new putative context genes.

**Figure 6:**
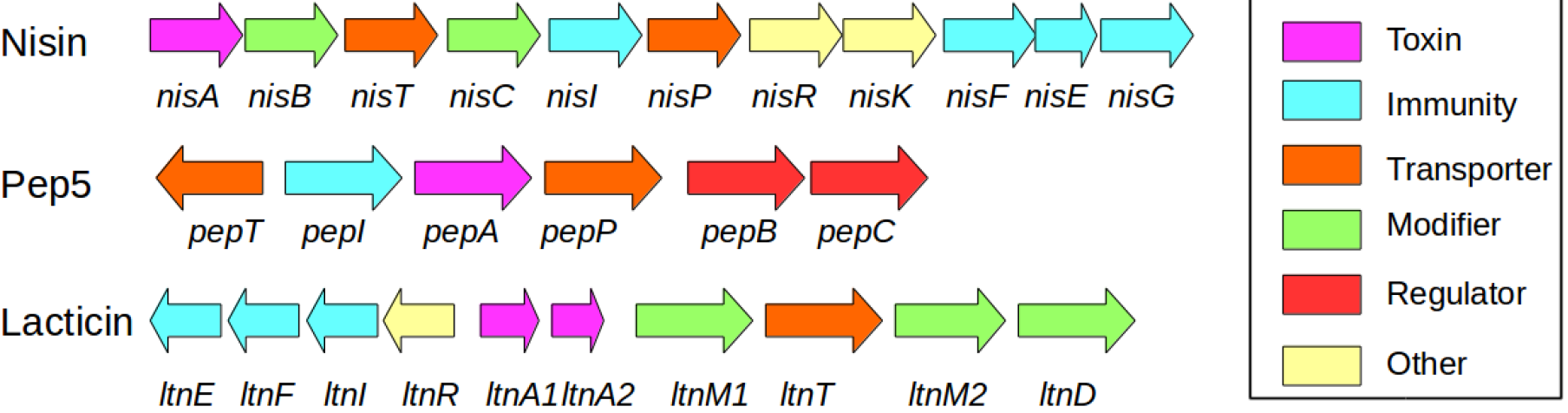
Bacteriocins with context genes. After[25].

We ran BLAST using all 1294 (54 experimentally verified and 1240 newly found) putative context genes against the whole bacteria RefSeq database [28] and we collected hits with an e-value ≤ 10^−6^. We separated all the hits by organism, arranged them by co-ordinates, and identified 50kb regions in the whole genome that have contiguous hits. We applied our trained machine learning model to predict putative bacteriocins in these 50kb regions.

### Datasets

We performed 10× cross-validations on the three datasets we built where the datasets consist of positive bacteriocins from BAGEL, and the three negative datasets we built from Uniprot Swissprot database.

The cross-validation itself was done 50 times with different random seeds for all cases except for the RNN, BLAST, and HMMER for which it was done 10 times due to computational time demand. For BLAST, a 35% sequence identity score was used as a threshold for calling a result positive. We used the same cross-validation folds for BLAST as other algorithms where we BLASTed the test set against the training set. For HMMER, an e-value of < 10^−3^ was used as the threshold for deciding if a sequence is bacteriocin. The reported results are the mean of 10× nested cross-validation done 50 times (10 times for RNN, BLAST and HMMER), and the standard error is from those 50 (10 for RNN, BLAST and HMMER) mean values.

## Results

Table S1 and Figure 7 show a comparison of Word2vec, trigram representation, BLAST and HMMER for the primary bacteriocin dataset in terms of precision, recall, and *F*_1_ score.

**Figure 7:**
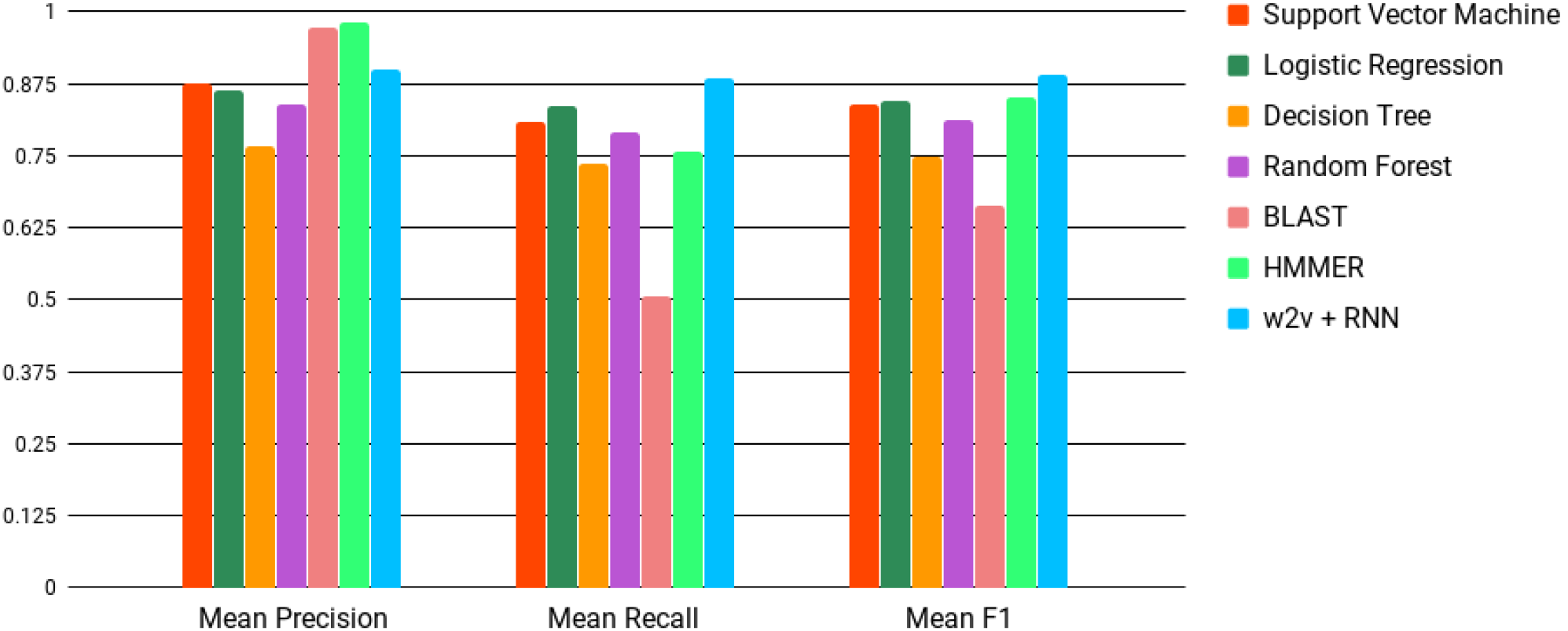
Mean *F*_1_ scores of different algorithms with both Word2vec (w2v) and baseline representation. Error bars are too small to show. W2v + RNN (blue) provides the best *F*_1_ score. See Table S1 for values and standard errors.

Precision (*Pr*) Recall (*Rc*) and *F*_1_ are defined as:

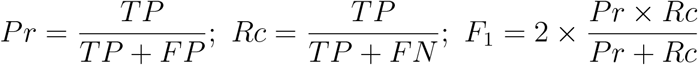

Where *TP*: True Positives, *FP*: False Positives, *FN*: False Negatives.

Recurrent Neural Network (RNN) with Word2vec representation for protein sequences provides the best recall, and *F*_1_ score. HMMER and BLAST have better precision scores which is expected as they only predict true positives by e-value and sequence identity respectively but they have high false negative rate. For trigram representation, support vector machine and logistic Regression perform similarly but with lower precision, recall, and *F*_1_ score than Word2vec with RNN. After getting the 10× cross-validation results, we trained the RNN on the whole data with the same hyper-parameters, and this final trained RNN was used to find new bacteriocins in the identified genomic regions.

Figure 8 shows the Precision-Recall curves for w2v+RNN, and for SVM, trigram+Logistic regression,trigram+ Random Forest, and BLAST. RNN has the largest area under the curve. BLAST is competitive with the other models except RNN. The curve for HMMER could not be shown as we need a probability value for each prediction which HMMER does not provide.

**Figure 8:**
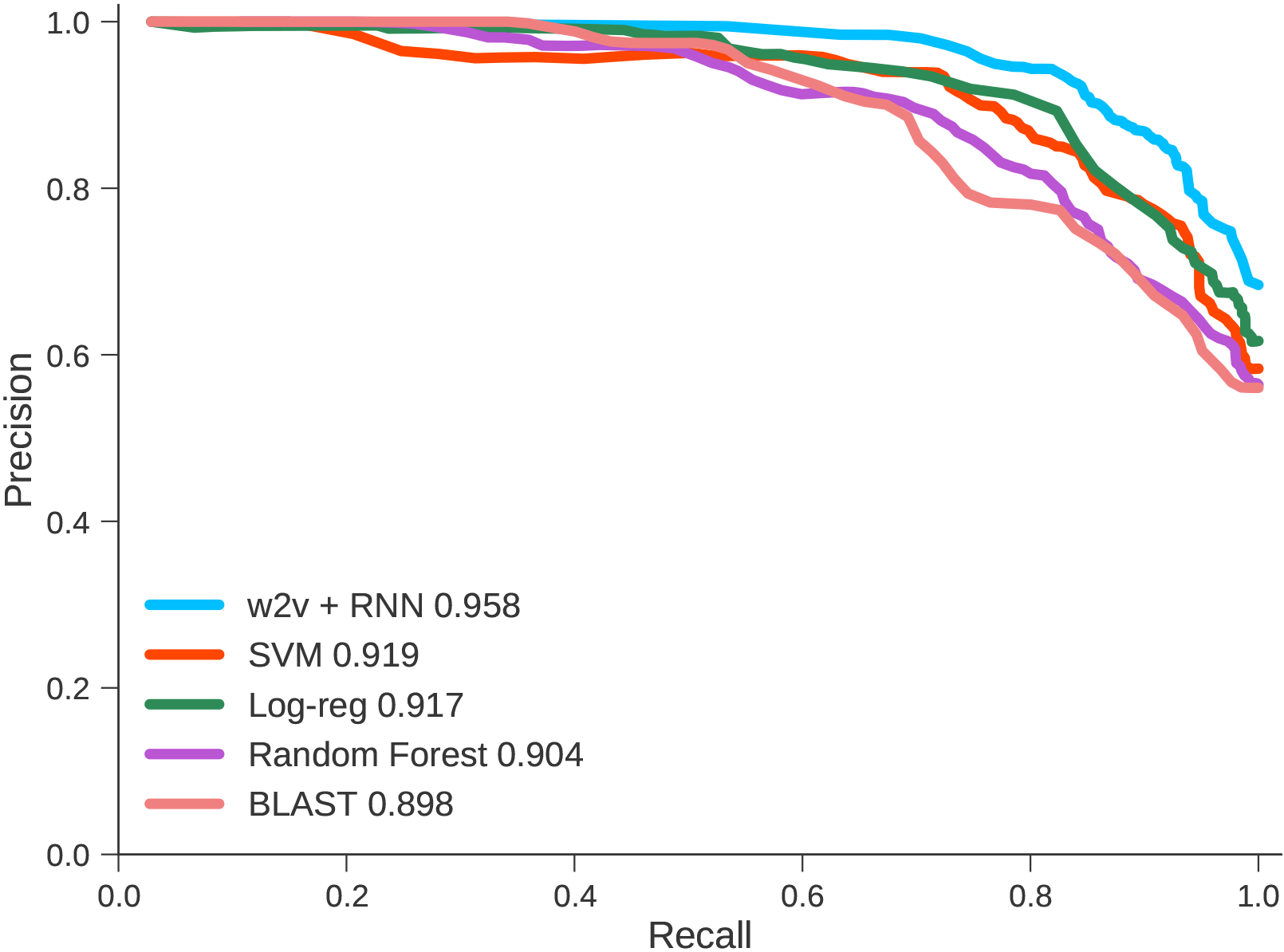
Mean Precision-Recall curves of one run of 10× cross validation for word2vec with RNN, Support Vector Machine (SVM), Logistic Regression (Log-reg), Random Forest, and BLAST. Number in legend is area under the curve. RNN performs better than all the other methods.

Table and S2 and S3 show the performance differences across the different bacteriocin data sets. he length distribution of the protein sequences in the positive and negative sets are different as mentioned in Methods. Looking at Table S2, The improvement in the w2v+RNN and the trigram based methods is evident, as well as the precison of HMMER. We assume the length disparity between the positive and negative sequences have helped in correctly classifying bacteriocins. Surprisingly, the precision of BLAST has decreased compared with its precision in the primary bacteriocin dataset. The performance of HMMER has largely remained the same over the different negative sets, its predictions more or less remain the same because of its low false positive rate. Table S3 shows the performance comparison for the third bacteriocin dataset. The length disparity between positive and negative sequences for the third dataset is even greater than the second bacteriocin dataset. SVM, Logistic Regression, Decision Tree, and Random Forest all have improved performance. SVM’s precision is comparable to that of w2v+RNN. RNN still has the best recall and *F*_1_ score. In contrast, BLAST’s performance has significantly decreased indicating that somehow the length disparity is causing problems in identifying true bacteriocins. Just like the second bacteriocin dataset, HMMER’s performance remains almost the same with a slight improvement on the precision score.

### Results on 50kb Chromosomal Stretches

We applied our trained w2v+RNN model on the sequences identified from the 50kb regions (see Methods) to predict putative bacteriocins. The w2v+RNN model predicted 119 putative bacteriocins with a probability of ≥ 0.99. Figure 9 shows three of our predicted bacteriocins in their genomic neighborhood *Lactobacillus*. We found several context genes surrounding these predicted bacteriocins, supporting our hypothesis that these bacteriocin predictions are valid.

**Figure 9.**
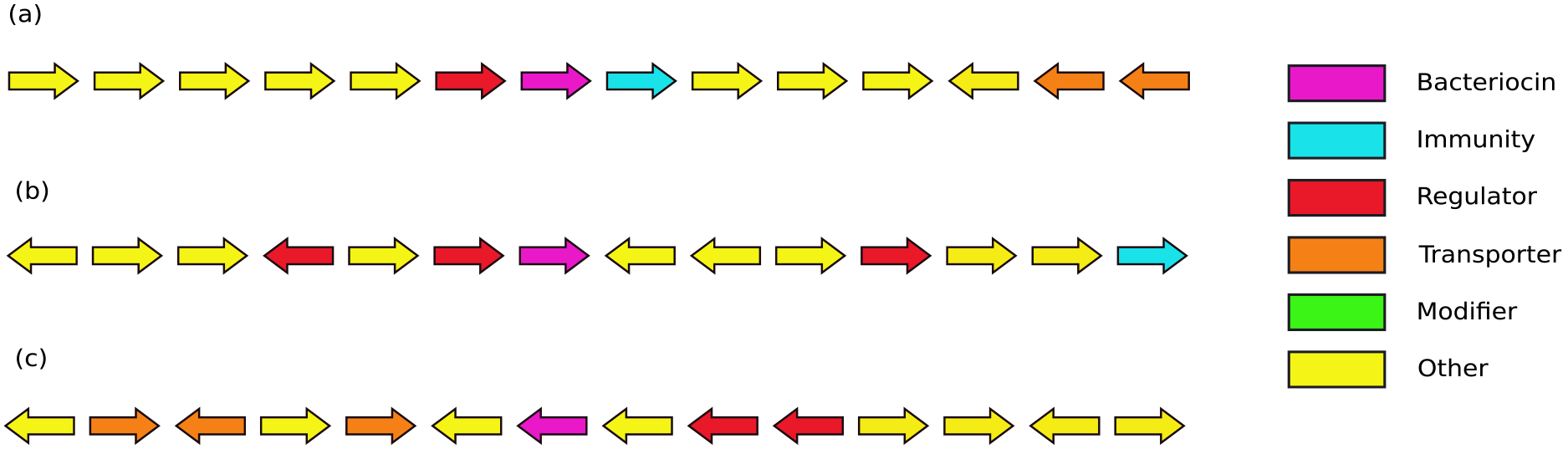
Context genes found surrounding the predicted bacteriocins within ±25kb range. (a) *Lactobacillus acidophilus NCFM* (Locus: NC_006814, putative bacteriocin: YP_193019.1, immunity: YP_193020.1, regulator: YP_193018.1, transporters: YP_193025.1, YP_193026.1) (b) *Lactobacillus helveticus R0052* (GenBnk: NC_018528, putative bacteriocin: YP_006656667.1, immunity: YP_006656674.1, regulator: YP_006656666.1, YP_006656664.1, YP_006656671.1) (c) *Lactobacillus helveticus CNRZ32* (GenBank ID: NC_021744, putative bacteriocin: YP_008236084.1, regulators: YP_008236086.1, YP_008236087.1, transporters: YP_008236082.1, YP_008236080.1, YP_008236079.1)

## Discussion

We developed a machine learning approach for predicting bacteriocins. Our approach does not require sequence similarity searches, and has discovered several putative bacteriocins with a high probability. The Word2vec representation takes advantage of the large volume of unlabeled bacterial protein sequences available. The Word2vec trigram representation can be used in other machine learning tasks in computational biology to represent protein sequences, for the purpose of discovering functional similarities cannot be discovered from sequence similarity. We used the embedding vectors for each overlapping trigram in a protein sequence, and used them as input in a temporal order for a RNN. Taken together, the embedding vectors with an RNN are used in natural language processing for common tasks such as sentence classification or document classification [29]. We also built three different datasets with three different negative bacteriocin sets. All the methods except RNN struggled to identify true bacteriocins in the primary bacteriocin dataset where the length distribution for positive and negative bacteriocins is exact. This is also the reason we used the primary bacteriocin dataset as the final dataset to train our RNN model before applying it to find novel bacteriocins in *Lactobacillus*. Compared with the primary bacteriocin dataset, the other methods except BLAST and HMMER have had improved performance as the differences in length distribution of positive and negative sequences increased in the second and third bacteriocin dataset. The pHMMs used (taken from the BOA [8] paper) for HMMER in our method were built using many sequences including the BAGEL dataset. Yet tested against the BAGEL sequences, its precision is high but the recall remained low compared with w2v+RNN.

Despite the training set being small, with proper regularization our RNN model provides better precision than all the other methods except BLAST and HMMER, and better recall than all other methods. We argue that embedding+RNN can be used to boost the prediction powers of machine learning models in sequence-based classification problems in biology. Our models also provide us with an associated confidence score, which is useful for experimentalists who wish to apply this method towards genome mining. We chose a threshold of 0.99 for RNN to provide the list of putative predictions. Although our training set is balanced in terms of bacteriocins and non-bacteriocins, the number of bacteriocin sequences in the microbial sequence universe is much lower. Finally, we provide six protein sequences that our (with probability of >= 0.99) model predicted to be putative bacteriocins where we could also find putative context genes. We also provide a set of total 119 sequences predicted by w2v+RNN with a probability of greater than 0.99. All of these sequences could not be detected against known bacteriocins when we used BLAST against the NR database with an e-value of 10^−3^ or less.

Historically, the use of bioinformatic prediction methods has favored high precision over high recall, as a large number of false positive findings can be costly for experiments that verify predictions. However, there are cases where it may be argued that a high recall method is appropriate. For example, with the need to cast a wider net in identifying potential drug candidates, coupled with the decrease in experimental costs. By employing a high recall method and choosing an appropriate accuracy threshold, experimentalists can calibrate the precision / recall tradeoff needed to optimize the functional testing of novel peptides.

Protein classification tasks are typically based on some form of sequence similarity as an indicator for evolutionary relatedness. However, in many cases non-orthologous replacements occur, where two non-homologous proteins perform the same function. Non-orthologous function replacements have been detected using natural language processing [30], genomic context methods [31, 32, 33], and other combined methods [34]. However, such methods require associated metadata or contextual genomic information. Here we present a solution to find functionally similar non-orthologs that does not require gathering these metadata, but does require a dataset of positive and negative examples. We therefore recommend that word embedding be explored for function classification involving dissimilar biological sequences.

We used the following software tools in this study: Keras [35], Scikit-learn [36], Gensim [37], Matplotlib [38], Jupyter notebooks [39], Numpy and Scipy [40].

## Availability

Data and code for this project are available at: https://github.com/nafizh/Bacteriocin_paper

## Funding

The research is based upon work supported, in part, by the Office of the Director of National Intelligence (ODNI), Intelligence Advanced Research Projects Activity (IARPA), via the Army Research Office (ARO) under cooperative Agreement Number W911NF-17-2-0105, and by the National Science Foundation (NSF) grant ABI-1551363. The views and conclusions contained herein are those of the authors and should not be interpreted as necessarily representing the official policies or endorsements, either expressed or implied, of the ODNI, IARPA, ARO, NSF, or the U.S. Government. The U.S. Government is authorized to reproduce and distribute reprints for Governmental purposes notwithstanding any copyright annotation thereon.

## Acknowledgements

Will be provided post-review.

